# Genome-wide association analysis identifies 27 novel loci associated with uterine leiomyomata revealing common genetic origins with endometriosis

**DOI:** 10.1101/324905

**Authors:** C. S. Gallagher, N. Mäkinen, H. R. Harris, O. Uimari, J. P. Cook, N. Shigesi, N. Rahmioglu, T. Ferreira, D. R. Velez-Edwards, T. L. Edwards, Z. Ruhioglu, F. Day, C. M. Becker, V. Karhunen, H. Martikainen, M-R Järvelin, R. M. Cantor, P. M. Ridker, K. L. Terry, J. E. Buring, S. D. Gordon, S. E. Medland, G. W. Montgomery, D. R. Nyholt, D. A. Hinds, J. Y. Tung, the 23andMe Research team, J. R. B. Perry, P. A. Lind, J. N. Painter, N. G. Martin, A. P. Morris, D. I. Chasman, S. Missmer, K. T. Zondervan, C. C. Morton

## Abstract

Uterine leiomyomata (UL), also known as uterine fibroids, are the most common neoplasms of the reproductive tract and the primary cause for hysterectomy, leading to considerable impact on women’s lives as well as high economic burden^1,2^. Genetic epidemiologic studies indicate that heritable risk factors contribute to UL pathogenesis^3^. Previous genome-wide association studies (GWAS) identified five loci associated with UL at genome-wide significance (*P* < 5 × 10^−8^)^4–6^. We conducted GWAS meta-analysis in 20,406 cases and 223,918 female controls of white European ancestry, identifying 24 genome-wide significant independent loci; 17 replicated in an unrelated cohort of 15,068 additional cases and 43,587 female controls. Aggregation of discovery and replication studies (35,474 cases and 267,505 female controls) revealed six additional significant loci. Interestingly, four of the 17 loci identified and replicated in these analyses have also been associated with risk for endometriosis – another common gynecologic disorder. These findings increase our understanding of the biological mechanisms underlying UL development, and suggest overlapping genetic origins with endometriosis.

---

UL are hormone-driven tumors that occur in 70-80% of all women by age 50^7^. Although the majority of UL are asymptomatic, about 25% of women with UL are symptomatic, and may experience excessive bleeding, abdominal pain, and infertility^2^. Currently, the only essentially curative treatment is uterine extirpation via total hysterectomy. Known risk factors for UL include increasing age up to menopause, ethnicity (particularly African ancestry), family history of UL, and increased body mass index (BMI)^3^. Studies on familial aggregation and twins, as well as racial differences in prevalence and morbidity, suggest heritable factors influence the risk for developing UL^8–13^. To date, previous GWAS including up to approximately 1,600 cases have identified five loci significantly associated (*P* < 5 × 10^−8^) with UL: 10q24.33, 11p15.5 and 22q13.1 in Japanese women^4^, 17q25.3 in white women of European ancestry^5^, and a distinct region at 22q13.1 in African American women^6^.

We conducted the largest discovery meta-analysis of GWAS on UL to date including four population-based cohorts of white European ancestry, increasing the case sample size almost 13-fold compared to previous studies (20,406 cases and 223,918 female controls): Women’s Genome Health Study (WGHS), Northern Finnish Birth Cohort (NFBC), QIMR Berghofer Medical Research Institute (QIMR), and The UK Biobank **(Supplementary Notes, Supplementary Table 1)**. Ancestry in each cohort was verified by principal component analysis (PCA) of genotype data. UL phenotype in each cohort was analyzed in a generalized linear regression model assuming additive genetic effects with multivariate adjustment for age and BMI, and correction for population structure. After quality control metrics were applied, including exclusion of non-informative (MAF < 0.01) and poorly imputed (*r*^2^ < 0.4) SNPs, we performed a fixed-effects, inverse-variance weighted meta-analysis. Altogether 8,292,290 biallelic SNPs were analyzed and adjustments for genomic inflation were performed **(Supplementary Fig. 1a, Supplementary Table 2)**. We identify a total of 1,167 SNPs across 24 loci with genome-wide significance (*P* < 5 × 10^−8^) **(Supplementary Table 3)**. The Manhattan plot is shown in **Fig. 1**.

**Fig. 1.**
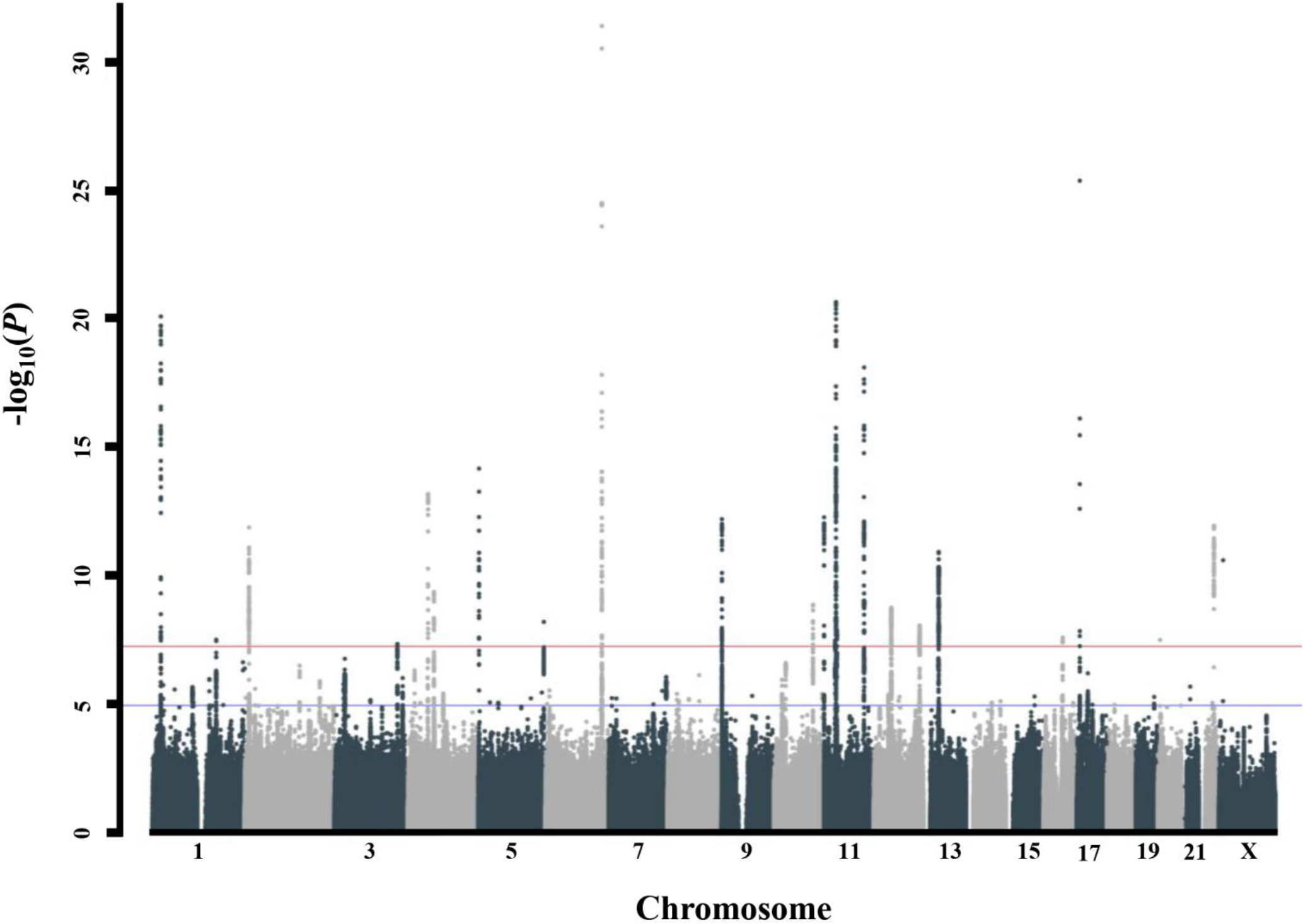
Manhattan plot for discovery-phase meta-analysis. Meta-analysis of GWAS across 244,324 women of white European ancestry conducted in population-based cohorts identifies 24 independent loci associated with UL. Red and blue horizontal lines indicate genome-wide significant (*P* < 5 × 10^−8^) and suggestive (*P* < 1 × 10^−5^) thresholds.

Replication of the peak SNPs was pursued in an independent direct-to-consumer cohort obtained from 23andMe, including 15,068 self-reported UL cases and 43,587 self-reported female controls of white European ancestry **(Supplementary Notes, Supplementary Fig. 1b, Supplementary Tables 1 and 2)**. We replicate associations at 17 of the 24 (71%) loci identified in our discovery-phase meta-analysis (significance threshold; *P* < 2.08 × 10^−3^) **(Table 1)**. Among independently replicated loci are all three loci previously reported to be associated with UL in Japanese women: 10q24.33 (rs9419958, odds ratio (OR) = 1.09, *P* = 1.20 × 10^−9^), 11p15.5 (rs547025, OR = 1.15, *P* with endometriosis: 1p36.12 (rs223552, OR = 1.14, *P* = 7.39 × 10^−21^), 2p25.1 (rs10929757, OR = 1.07, *P* = 1.19 × 10^−12^), 6q25.2 (rs58415480, OR = 1.18, *P* = 3.65 × 10^−32^), and 11p14.1 = 4.77 × 10^−13^), and 22q13.1 (rs12484776, OR = 1.09, *P* = 4.08 × 10^−12^)^4^.

**Table 1.**
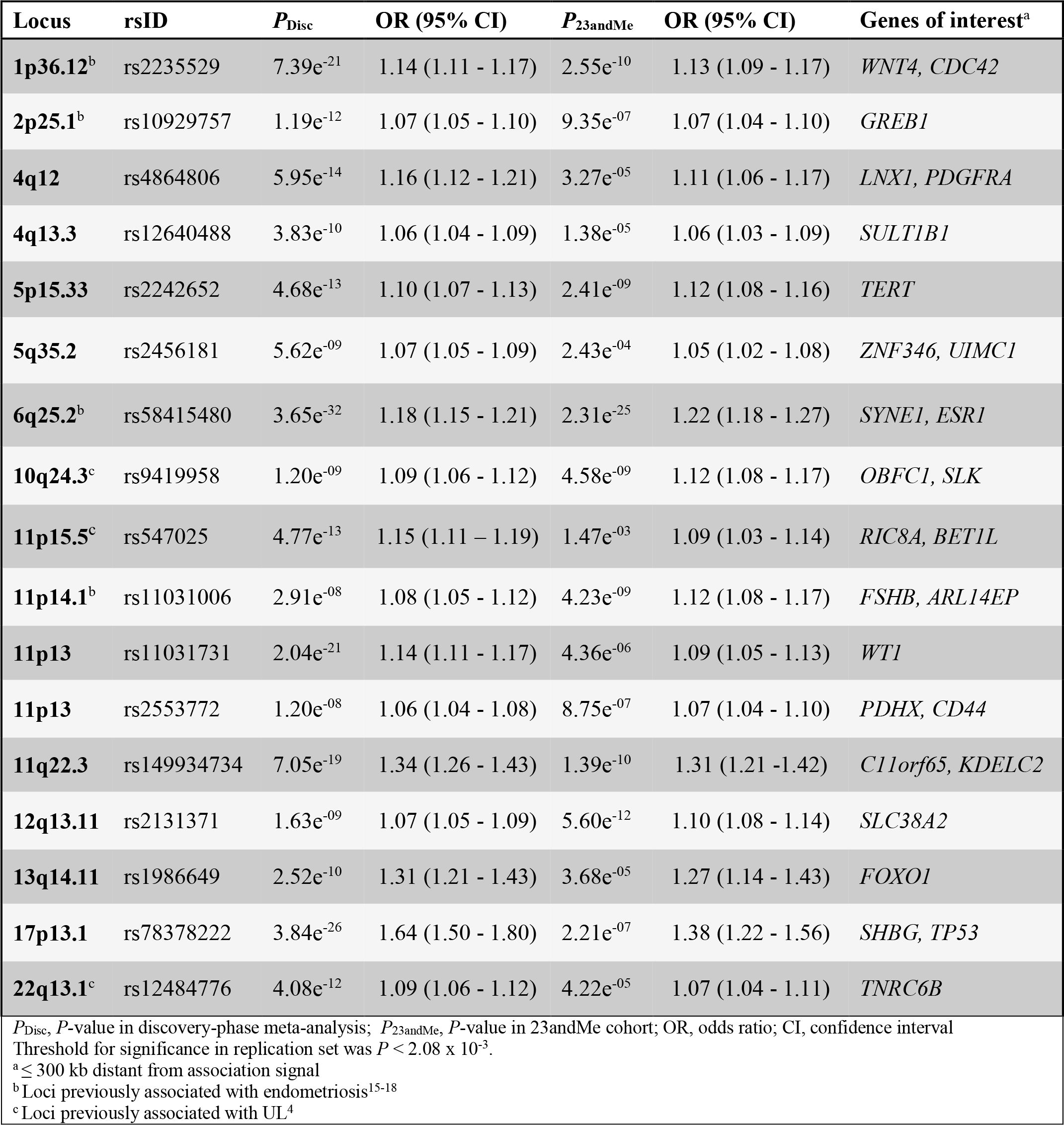
Seventeen loci identified in discovery-phase GWAS meta-analysis and replicated in an independent cohort from 23andMe.

Interestingly, significant association signals are also observed at several loci previously associated with endometriosis: 1p36.12 (rs223552, OR = 1.14, *P* = 7.39 × 10^−21^), 2p25.1 (rs10929757, OR = 1.07, *P* = 1.19 × 10^−12^), 6q25.2 (rs58415480, OR = 1.18, *P* = 3.65 × 10^−32^), and 11p14.1 (rs11031006, OR = 1.08, *P* = 2.91 × 10^−8^)^14–17^. Endometriosis is another common hormone-dependent disease that affects reproductive-aged women, resulting from ectopic growth of endometrial tissue outside the uterine cavity^18^. Although functional studies of relevant tissue need to confirm the consequences of the variants in regulation of gene expression, each of the overlapping genomic loci contain a gene(s) known to be involved in progesterone or estrogen signaling. *WNT4* at 1p36.12 encodes a secreted signaling factor that promotes female sex development, and regulates both postnatal uterine development and progesterone signaling during decidualization^19,20^. Recently, SNPs increasing endometriosis risk at 1p36.12 have been suggested to act through *CDC42*, a gene which encodes a small GTPase of the Rho family ^21^. *GREB1* at 2p25.1 is an early response gene in the estrogen receptor (ER)-regulated pathway, and promotes growth of breast and pancreatic cancer cells^22,23^. *ESR1* at 6q25.2 encodes the alpha subunit of the ligand-activated nuclear ER that regulates cell proliferation in the uterus^24^. *FSHB* at 11p14.1 encodes the biologically active subunit of follicle-stimulating hormone, which regulates maturation of ovarian follicles and release of ova during menstruation^25,26^.

A number of replicated loci also harbor genes previously implicated in cell growth and cancer risk in different tissue types, including cervical cancer^27^, epithelial ovarian cancer^28,29^, breast cancer^30,31^, glioma^32,33^, bladder cancer^34^, and pancreatic cancer^35–37^. Specifically, seven of the independent loci contain well-characterized oncogenes and tumor suppressor genes from the Cancer Gene Census list in COSMIC^38^: *PDGFRA*, *TERT, ESR1*, *WT1*, *ATM*, *FOXO1*, and *TP53*. FOXO1 is a transcription factor that plays an important role in cell proliferation, apoptosis, DNA repair, and stress response^39^. Inactivation of *FOXO1* promotes cell proliferation and tumorigenesis in several hormone-regulated malignancies, such as prostate, breast, cervical, and endometrial cancers^40–43^. We quantified and compared nuclear FOXO1 expression in 335 UL and 35 patient-matched normal myometrial samples using immunohistochemistry on tissue microarrays. In the patient-matched tumor-normal pairs, nuclear FOXO1 expression was 1.69-fold higher (*P* = 0.01) in UL **(Supplementary Fig. 2a)**. When all 335 UL were taken into account, expression was increased as much as 2.32-fold (*P* = 1.52 × 10^−9^) in UL compared to myometrial samples (**Supplementary Fig. 2b**). A total of 109 patient samples were genotyped for rs6563799 and rs7986407 – two of the peak SNPs residing in the *FOXO1* locus. Stratification of samples by genotype reveals a statistically significant increase in FOXO1 levels of UL harboring the risk allele for rs6563799 (allelic dosage, *P* = 0.047; homozygosity for risk allele, *P* = 0.035) **(Supplementary Fig. 3a**). An increase in FOXO1 levels of UL with the rs7986407 risk allele is also observed; however, the change is not statistically significant **(Supplementary Fig. 3b)**. Our results are consistent with a previous study reporting elevated FOXO1 expression in UL compared to matched myometrium^44^, especially levels of phosphorylated (p) FOXO1 (pSer^256^). Normally, p-FOXO1 interacts with 14-3-3γ protein in the nucleus, resulting in translocation of FOXO1 to the cytoplasm and its transcriptional inactivation. Interestingly, Kovacs et al. showed p-FOXO1 to be predominantly present in the nucleus in UL, but sequestered in the cytoplasm of myometrium^44^. The concomitant increase of p-FOXO1 and reduced expression of 14-3-3γ in UL has been suggested to lead to impaired nuclear/cytoplasmic shuttling of p-FOXO1, which promotes cell survival^44–46^.

Next, we conducted a meta-analysis on a total of 8,602,260 SNPs across all cohorts, including a total of 35,474 UL cases and 267,505 controls **(Supplementary Fig. 1c, Supplementary Table 2)**. Through linkage disequilibrium score (LDSC) regression analysis, an estimated 89.5% of the genomic inflation factor (λGC) of 1.12 was attributable to polygenic heritability (intercept = 1.02, s.e. = 0.0081). We observe genome-wide significant associations (*P* < 5 × 10^−8^) at 2,045 SNPs across 27 independent loci **(Supplementary Fig. 4, Supplementary Table 4)**. The Manhattan plot is shown in **Fig. 2**. In addition to 21 loci identified at the discovery stage, we observe six novel loci significantly associated with UL **(Table 2)**. Among the novel, ‘discovery’ loci are three genes of interest: *HMGA1*, *BABAM2*, and *WNT2*. *HMGA1* is a member of the high mobility group proteins and is involved in regulation of gene transcription^47^. Somatic rearrangements of *HMGA1* at 6p21 have been recurrently documented in UL, albeit at a much lower frequency than those of *HMGA2* – another member of the protein family^48–50^. *BABAM2* encodes a death receptor-associating intracellular protein that promotes tumor growth by suppressing apoptosis^51^. Associations at the locus containing *WNT2* together with associations observed at the *WNT4* locus reinforce a possible role for Wnt signaling in UL. Overall, individual SNP-based heritability (*h*^*2*^) was estimated to be relatively low, 0.0281 (s.e. = 0.0029) on the liability scale.

**Fig. 2.**
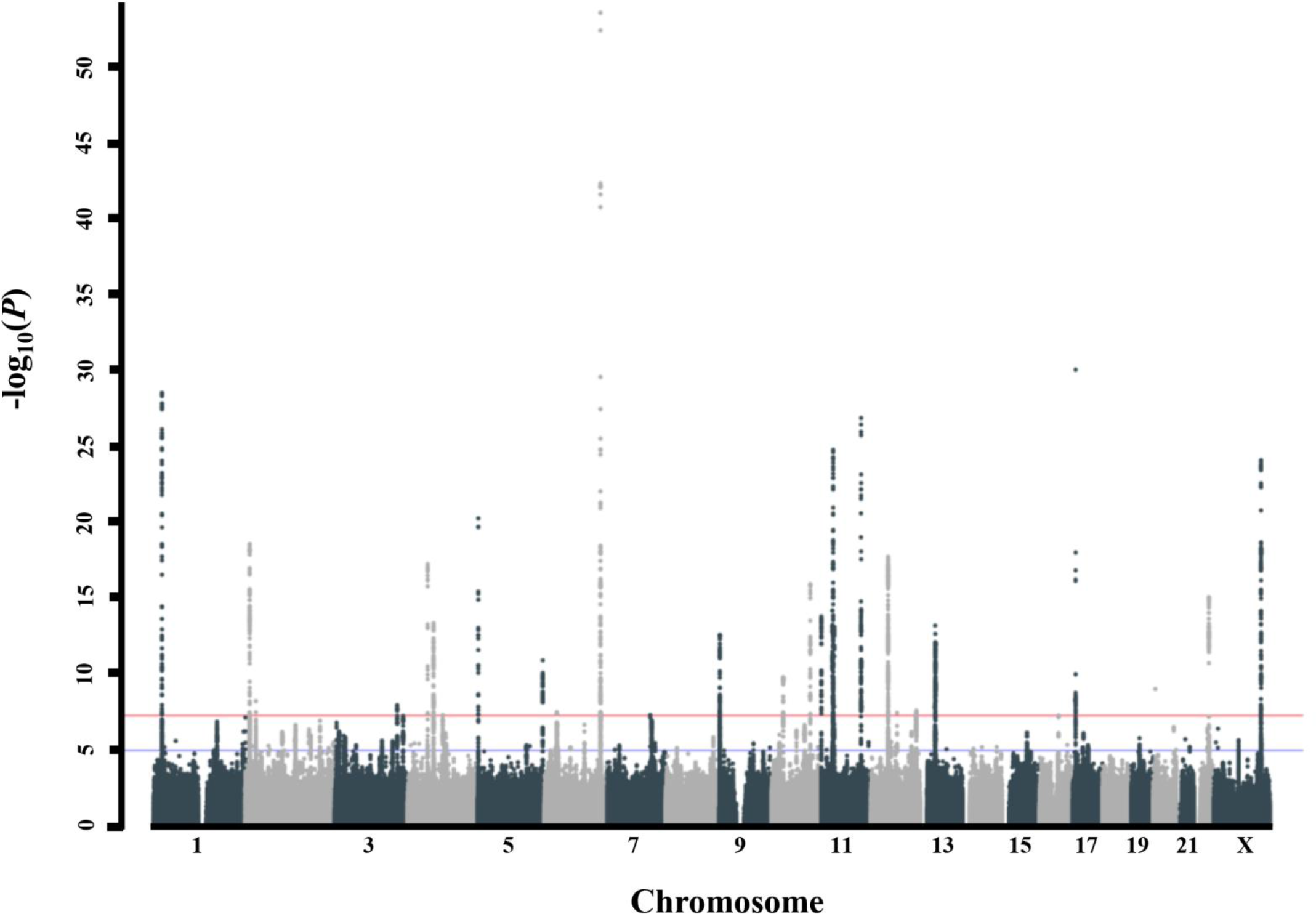
Manhattan plot for meta-analysis across all cohorts from the FibroGENE consortium. Meta-analysis of GWAS across 302,979 women of white European ancestry across all cohorts identifies 27 independent loci associated with UL. Red and blue horizontal lines indicate genome-wide significant (*P* < 5 × 10^−8^) and suggestive (*P* < 1 × 10^−5^) thresholds.

**Table 2.** Six novel genome-wide significant loci identified in meta-analysis across all cohorts.

| **Locus** | **rsID** | **$P_{\text{Meta}}$** | **OR (95% CI)** | **Genes of interest<sup>a</sup>** |
| --- | --- | --- | --- | --- |
| 2p23.2 | rs55819434 | 5.59e<sup>-09</sup> | 1.09 (1.06 – 1.12) | *BABAM2* |
| 4q22.3 | rs4699299 | 4.72e<sup>-08</sup> | 1.05 (1.03 – 1.07) | *PDLIM5* |
| 6p21.31 | rs116251328 | 2.95e<sup>-08</sup> | 1.15 (1.09 – 1.21) | *GRM4, HMGA1* |
| 7q31.2 | rs2270206 | 4.64e<sup>-08</sup> | 1.06 (1.04 – 1.09) | *WNT2* |
| 10p11.22 | rs10508765 | 1.51e<sup>-10</sup> | 1.07 (1.05 – 1.09) | *ZEB1, ARHGAP12* |
| 12q15 | rs11178393 | 3.34e<sup>-08</sup> | 1.08 (1.05 – 1.10) | *PTPRR* |
| OR, odds ratio; CI, confidence interval<br><sup>a</sup> ≤ 300 kb distant from association signal |  |  |  |  |

Gene-set and tissue enrichment analyses across 5,185 SNPs with suggestive (*P* < 1 × 10^−5^) or significant (*P* < 5 × 10^−8^) UL associations using DEPICT^52^ reveal significant enrichments (false discovery rate (FDR) < 0.05) in gene sets, such as steroid hormone receptor (GO:0035258; *P* = 8.09 × 10^−6^), hormone receptor binding (GO:0051427; *P* = 1.49 × 10^−4^), and nuclear hormone receptor binding (GO:0035257; *P* = 9.71 × 10^−5^) **(Supplementary Tables 5 and 6)**. The results are concordant with the hormone-driven nature of UL. We also observe enrichment of genes associated with expression in female urogenital tissue (*P* = 6.19 × 10^−4^) **(Supplementary Fig. 5)**. To identify SNPs with likely regulatory function, we selected up to 30 of the most significant SNPs from each of the 27 loci identified in the meta-analysis of GWAS across all cohorts. Altogether 429 of 597 SNPs (72%) were present in the RegulomeDB^53^, and 23 of these have a score < 3, indicating potential involvement in gene regulation **(Supplementary Table 7)**. Based on the RegulomeDB, two SNPs (rs498217 at 11p15.5 and rs1641528 at 17p13.1) are indicated as potential expression quantitative trait loci (eQTLs) in monocytes for *SCGB1C1* and *CD68*, respectively^54^.

In summary, our GWA analyses uncovered 27 novel genomic loci associated with UL in women of white European ancestry. Many of the candidates fall into two categories: (1) genomic regions containing characterized tumor suppressors and oncogenes, and (2) genes involved in hormone signaling pathways previously associated with endometriosis. Biological overlap between two highly common gynecologic diseases, due to similarities in molecular mechanisms and progenitor cells, has long been suspected. Further characterization of the mutual pathogenic mechanisms has the capacity to direct not only a deeper understanding of the underlying biology, but also treatments for two diseases that cause significant morbidity in roughly one-third of the world’s population.

## Online Methods

### Subjects

Four population-based cohorts (WGHS, NFBC, QIMR, and UK Biobank) and one direct-to-consumer cohort (23andMe) from the FibroGENE consortium were included in the study **(Supplementary Table 1)**, resulting in 35,474 UL cases and 267,505 female controls of white European ancestry. Detailed descriptions of patient cohorts and sample selection are available in **Supplementary Notes**. For the current study, sample sizes were maximized using a basic, harmonizing phenotype definition to separate cases and controls solely based on either self-report or clinically documented UL history. All participants provided an informed consent in accordance with the processes approved by the relevant jurisdiction for human subject research for each cohort.

### Genotyping

For the GWAS discovery stage, several different Illumina-based genotyping platforms (Illumina Inc., San Diego, CA, USA) were used: HumanHap300 Duo‘+’ chips or the combination of the Human-Hap300 Duo and iSelect chips (WGHS), Infinium 370cnvDuo array (NFBC), 317K, 370K, or 610K SNP platforms (QIMR). Genotyping of participants in the UK Biobank was performed either on the Affymetrix UK BiLEVE or Affymetrix UK Biobank Axiom^®^ array with over 95% similarity. Genotyping of participants in the 23andMe cohort was performed on various versions of Illumina-based BeadChips.

### Quality control and imputation

Each cohort conducted quality control measures and imputation for their data. For WGHS, NFBC, QIMR, and 23andMe, all cases and controls with a genotyping call rate < 0.98 were excluded from the study. Imputation was performed on both autosomal and sex chromosomes using the reference panel from the 1000 Genomes Project European dataset (1000G EUR) Phase 3. Imputation was carried out using ShapeIt2 and IMPUTE2 softwares^55,56^. SNPs with call rates of < 99%, and SNPs showing deviation from Hardy-Weinberg equilibrium (*P* ≤ 1 × 10^−6^) were excluded from further analyses. Population-stratification for the data was examined with principal component analysis (PCA) using EIGENSTRAT^57^. The four HapMap populations were used as reference groups: Europeans (CEU), Africans (YRI), Japanese (JPT), and Chinese (CHB). All observed outliers were removed from the study. The UK Biobank data QC and imputation was handled by a dedicated team headed by the Wellcome Centre for Human Genetics, prior to public release of the data. Genotype data used in the present analyses were imputed up to the Haplotype Reference Consortium (HRC) panel. We applied additional quality control filters to exclude poorly imputed SNPs (*r*^2^ < 0.4) and SNPs with a MAF of < 1%.

### Association analyses

Using additive encoding of genotypes and adjusting for age, BMI, and the first five principal components from EIGENSTRAT, WGHS, NFBC, QIMR, and 23andMe cohorts in both the discovery and replication stage performed logistic regression analysis and provided summary statistics, including beta coefficients, χ^2^ values, and standard errors, for the genotyped and imputed SNPs. The UK Biobank association analyses were conducted using a linear mixed model (BOLT-LMM v.2.3.2)^58^ adjusting for the two array types used, age and BMI (fixed effects) and a random effect adjusting for relatedness between women. Effect size estimates (β and SE) from the linear mixed-model were converted to log-odds scale prior to meta-analysis. Two fixed-effects, inverse variance weighted meta-analyses on summary statistics were conducted using METAL^59^, one in the discovery stage and the other across all cohorts. For discovery-phase meta-analysis, 8,292,290 biallelic SNPs for which data were available in at least two of the four cohorts were analyzed. A total of 8,602,260 SNPs were available from at least two of the five cohorts for the meta-analysis across all cohorts. Quantile-quantile plots of the results from meta-analysis of population-based cohorts and across all GWAS cohorts are shown in **Supplementary Fig. 1**. Details on the overall genomic inflation factor and number of analyzed SNPs for each cohort are provided in **Supplementary Table 2.** Independence of genetic association with UL was defined as SNPs in low linkage disequilibrium (LD; *r*^2^ < 0.1) with nearby (≤ 500kb) significantly associated SNPs. Individual loci correspond to regions of the genome containing all SNPs in LD (*r*^2^ > 0.6) with index SNPs. Any adjacent regions within 250 kb of one another were combined and classified as a single locus of association.

### FOXO1 immunohistochemistry and genotyping

FOXO1 immunostaining was performed on two replicate tissue microarrays (TMAs) containing 335 UL and 35 patient-matched myometrium tissue samples from 200 white women of European ancestry obtained from myomectomies and hysterectomies. Tissue cores on the replicate TMAs represent different regions of the same samples, which include corresponding tumor-normal tissue pairs from 34 women. Immunohistochemistry was carried out using the BOND staining system (Leica Biosystems, Buffalo Grove, IL) with a primary antibody dilution of 1:100 (clone C29H4, Cell Signaling Technology, Danvers, MA) and hematoxylin as the counterstain. Immunostaining was analyzed using Aperio ImageScope software (Leica Biosystems). Each core was evaluated for the ratio of stain to counterstain taking into account variable cellularity between cores. Only nuclear labeling of the protein was evaluated. The average stain-to-counterstain ratio was compared between patient-matched UL and myometrium samples using a paired *t*-test (two-tailed), while an unpaired *t*-test (Welch’s *t*-test, two-tailed) was applied to compare all UL and myometrium samples. Genomic DNA from 109 UL on the TMA was available for genotyping. These UL were genotyped for two SNPs with genome-wide significance at the 13q14.11 locus: rs6563799 and rs7986407. For each SNP, the average FOXO1 stain-to-counterstain ratio was compared across increasing dosage of the risk allele using a one-way analysis of variance test (two-tailed). We also performed an unpaired *t*-test to compare mean expression of UL homozygous for the risk variant against the other genotypes (Welch’s *t*-test, two-tailed). *P*-values < 0.05 were considered statistically significant.

### Linkage disequilibrium score regression (LDSC)

Analysis of residual inflation in test statistics was conducted using univariate LDSC regression. Individual χ^2^ values for each SNP analyzed in the GWAS meta-analysis were regressed onto LD scores estimated from the 1000G EUR panel. Heritability calculations can be derived from analyzing the slope and y-axis intercept of the slope of the regression line. Percent impact of confounders, such as population stratification, on test statistic inflation are quantified as the LDSC ratio [((intercept – 1)) / ((mean χ^2^ – 1))] * 100%. Remaining effects [(1 – LDSC ratio) * 100%] represent the percentage of inflation attributed to polygenic heritability. Univariate LDSC regression was conducted using the LDSC software (https://github.com/bulik/ldsc.git). Adjustment of heritability (*h*^2^) calculations to the liability scale were performed by accounting for the prevalence of UL in the sample (~0.132) compared to the general population (~0.300).

### Gene-set and tissue enrichment analyses

Summary statistics from the set of 5,185 SNPs with suggestive (*P* < 1 × 10^−5^) or significant associations (*P* < 5 × 10^−8^) were analyzed for gene-set and tissue enrichment using the Data-driven Expression-Prioritized Integration for Complex Traits (DEPICT) software^52^. Using the 1000G EUR panel as a reference for LD calculations and the ‘clumping’ algorithm in PLINK^60^, we identified 162 independent loci at the suggestive threshold for DEPICT analyses **(Supplementary Table 5)**. FDR < 0.05 was considered statistically significant.

### URLs

WHS, http://whs.bwh.harvard.edu/; NFBC, http://www.oulu.fi/nfbc/; QIMR, http://www.qimrberghofer.edu.au/; UK Biobank, http://www.ukbiobank.ac.uk/; 23andMe, https://research.23andme.com/; METAL, http://csg.sph.umich.edu/abecasis/metal/; LDSC, https://github.com/bulik/ldsc.git; DEPICT, https://data.broadinstitute.org/mpg/depict/; RegulomeDB, http://www.regulomedb.org/; PLINK, http://pngu.mgh.harvard.edu/purcell/plink/

## Supplementary Material

Supplementary Material includes Supplementary Notes, five figures, and seven tables.

## Acknowledgements

The authors thank all of the women and their families who participated in WGHS, NFBC, QIMR, UK Biobank, and 23andMe. This study was supported by National Institutes of Health (NIH)/Eunice Kennedy Shriver National Institute of Child Health and Human Development (NICHD) grant (R01HD060530). We thank the Dana-Farber/Harvard Cancer Center in Boston, MA, for the use of the Specialized Histopathology Core, which provided FOXO1 immunostaining service. The Dana-Farber/Harvard Cancer Center is supported in part by an NCI Cancer Center Support Grant # NIH 5 P30 CA06516. **WGHS:** WGHS is supported by the National Heart, Lung, and Blood Institute (HL043851 and HL080467) and the National Cancer Institute (CA047988 and UM1CA182913) with funding for genotyping provided by Amgen. **NFBC:** NFBC1966 received financial support related to this study from the Academy of Finland (project grants 104781, 120315, 129269, 1114194, 24300796, 85547, Center of Excellence in Complex Disease Genetics), University Hospital Oulu, Biocenter, University of Oulu, Finland (75617), NHLBI grant 5R01HL087679-02 through the STAMPEED program (1RL1MH083268-01), NIH/NIMH (5R01MH63706:02), the EU FP5 EURO-BLCS, QLG1-CT-2000-01643, ENGAGE project and grant agreement HEALTH-F4-2007-201413, EU FP7 EurHEALTHAgeing 277849, the Medical Research Council (MRC), UK (G0500539, G0600705, G1002319, PrevMetSyn/SALVE) and ERDF European Regional Development Fund Grant no. 539/2010 A31592. The program is currently being funded by the EU H2020--PHC-2014 DynaHEALTH action (grant agreements No. 633595), EU H2020-HCO-2004 iHEALTH Action, EU H2020-PHC-2014 ALEC Action (grant agreement No. 633212), EU H2020-SC1-2016-2017 LIFECYCLE Action, EU H2020-MSCA-ITN-2016 CAPICE Action, Academy of Finland EGEA-project (285547) and MRC Grant MR/M013138/1. DNA extractions, sample quality controls, biobank up-keeping, and aliquotting were performed in the National Public Health Institute, Biomedicum Helsinki, Finland and supported financially by the Academy of Finland and Biocentrum Helsinki. We thank the late Professor Paula Rantakallio for the launch of NFBC1966. For further information, contact Professor Marjo-Riitta Jarvelin. **QIMR:** Funding for the twin studies was provided by the Australian National Health and Medical Research Council (241944, 339462, 389927, 389875, 389891, 389892, 389938, 442915, 442981, 496739, 552485, 552498, 1084325), the Australian Research Council (A7960034, A79906588, A79801419, DP0770096, DP0212016, DP0343921), the FP-5 GenomEUtwin Project (QLG2-CT-2002-01254), and the U.S. National Institutes of Health (NIH grants AA07535, AA10248, AA13320, AA13321, AA13326, AA14041, MH66206). The endometriosis study was supported by grants from the Australian National Health and Medical Research Council (241944, 339462, 389927,389875, 389891, 389892, 389938, 443036, 442915, 442981, 496610, 496739, 552485, 552498), the Cooperative Research Centre for Discovery of Genes for Common Human Diseases (CRC), Cerylid Biosciences (Melbourne), and donations from Neville and Shirley Hawkins. **UK Biobank:** This research has been conducted using the UK Biobank Resource under application 9637. This work was supported by the Medical Research Council [Unit Programme number MC_UU_12015/2]. **23andMe:** Collaborators for the 23andMe Research Team are: Michelle Agee, Babak Alipanahi, Adam Auton, Robert K. Bell, Katarzyna Bryc, Sarah L. Elson, Pierre Fontanillas, Nicholas A. Furlotte, Karen E. Huber, Aaron Kleinman, Nadia K. Litterman, Matthew H. McIntyre, Joanna L. Mountain, Elizabeth S. Noblin, Carrie A.M. Northover, Steven J. Pitts, J. Fah Sathirapongsasuti, Olga V. Sazonova, Janie F. Shelton, Suyash Shringarpure, Chao Tian, Vladimir Vacic, Catherine H. Wilson. **Personal grants:** N.M. acknowledges support from the Academy of Finland (295693). H.R.H. is supported by NIH K22 CA193860. T.F. is supported by the NIHR Biomedical Research Centre, Oxford. S.E.M. is supported by the National Health and Medical Research Council (NHMRC) Fellowship Scheme (1103623).

